# PAMalytics: a no-code application for structured validation of bioacoustic detections

**DOI:** 10.64898/2026.08.14.744822

**Authors:** Alastair Pickering, Santiago Martinez Balvanera, Nicholas Brown, Sareach Chea, Rhys Preston-Allen, Ratha Sor, Daniel S. Maynard, Jenna Lawson

## Abstract

1. Passive acoustic monitoring (PAM) is increasingly used for ecological research, biodiversity monitoring, assessment, and reporting. Automated species classifiers make it feasible to process large audio datasets but generate numerous detections that often need validation before use in downstream analyses or formal outputs.

2. Method development in PAM has focused on classifier building and downstream models that account for imperfect detection, yet the practical step between these - post-classification validation - remains weakly supported and is often implemented through ad hoc workflows. This increases manual handling, creates scope for transcription or consolidation errors, limits transparency and makes it difficult to document what was reviewed.

3. We introduce PAMalytics, an open-source, no-code, local browser-based application to support post-classification validation as a standardised workflow stage. PAMalytics ingests detections from any classifier, allows users to define how detections are sampled for review, and presents selected detections alongside their spectrograms with audio playback in one unified interface. Sampling strategy and review decisions are tracked alongside reviewer identity improving traceability and reproducibility across the validation workflow.

4. Case studies with Conservation International Cambodia and Imperial College London demonstrate PAMalytics in two validation settings. In Cambodia, gibbon predictions from a large, uneven dataset were sampled within sites, with likely classifier errors prioritised for validation. At Imperial, Amazon bird detections were sampled across each species’ classifier-confidence range before biodiversity metrics were derived. In both cases, PAMalytics reduced manual handling and validation time. By turning an ad hoc step into an accessible, structured workflow for conservation practitioners, PAMalytics fills a practical gap in the PAM bioacoustics pipeline and strengthens the link between automated detections and evidence used in biodiversity monitoring and reporting.

## 1 INTRODUCTION

Passive acoustic monitoring (PAM) – the systematic recording and analysis of biological sounds in the environment – has become a powerful tool for ecological research and conservation practice in recent decades (Stowell, 2022). Inexpensive and robust recording devices, together with expanding storage and processing capacity, now make it possible to collect terabytes of audio from remote and biodiverse landscapes, often over months or years of deployment (Gibb *et al*., 2019; Sugai *et al*., 2019). In parallel, automated bioacoustic classifiers, such as BirdNET (Kahl *et al*., 2021) and BatDetect2 (Aodha *et al*., 2022), have dramatically reduced the amount of manual effort required to work with these datasets by automatically assigning putative species labels and associated confidence scores to very large numbers of audio files.

As these tools have matured, PAM has increasingly moved into operational conservation and environmental management workflows (Aide *et al*., 2013; Browning *et al*., 2017). Government agencies, non-governmental organisations and local communities now use autonomous recorders alongside trained classifiers to monitor threatened species, evaluate land-use and restoration interventions, and provide evidence for regulatory compliance (Marques *et al*., 2013; Browning *et al*., 2017; NatureScot, 2021). As a result, outputs from automated classifiers increasingly feed into formal assessment, reporting and accountability processes, including environmental impact assessment (EIA), compliance monitoring and corporate or institutional biodiversity reporting (Cord *et al*., 2025; Miller *et al*., 2025).

While automated bioacoustic classifiers have largely alleviated the original scalability constraint in PAM - the need to manually inspect large volumes of audio (Gibb *et al*., 2019; Stowell, 2022) - they have shifted the practitioner burden downstream. Large volumes of classifier detections still often need to be validated, typically through targeted or subsampled review, before they can be used for analysis and reporting (Cowans *et al*., 2024; Malerba *et al*., 2026). Here, validation is defined as a quality-assurance step in which a defined subset of detections is checked manually by trained human reviewers and necessary adjustments are recorded so that downstream reporting can be accompanied by true error estimates. Practitioners can use established software to run detectors or classifiers, visualise and annotate sounds, and manage larger acoustic datasets and workflows, including Kaleidoscope, PAMGuard, Raven, Whombat and Arbimon (Gillespie *et al*., 2009; Aide *et al*., 2013; Martínez Balvanera *et al*., 2025). While some of these tools feature validation interfaces, they are often tied to specific classifiers or require cloud dependencies. Downstream, substantial work has addressed how to account for imperfect detection, especially through occupancy and related hierarchical models that allow for false negatives and false positives (Chambert *et al*., 2018; Barré *et al*., 2019; López-Baucells *et al*., 2019; Wright *et al*., 2020; Katsis *et al*., 2025). Between these stages, however, the workflow that links automated detections to ecological inference is comparatively unstructured (Fig. 1), despite the growing scale of classifier outputs and the recognised need for this step in applied analyses (Chambert, Miller and Nichols, 2015; Chambert *et al*., 2018; Metcalf *et al*., 2022; Cowans *et al*., 2024; Malerba *et al*., 2026).

**Figure 1:**
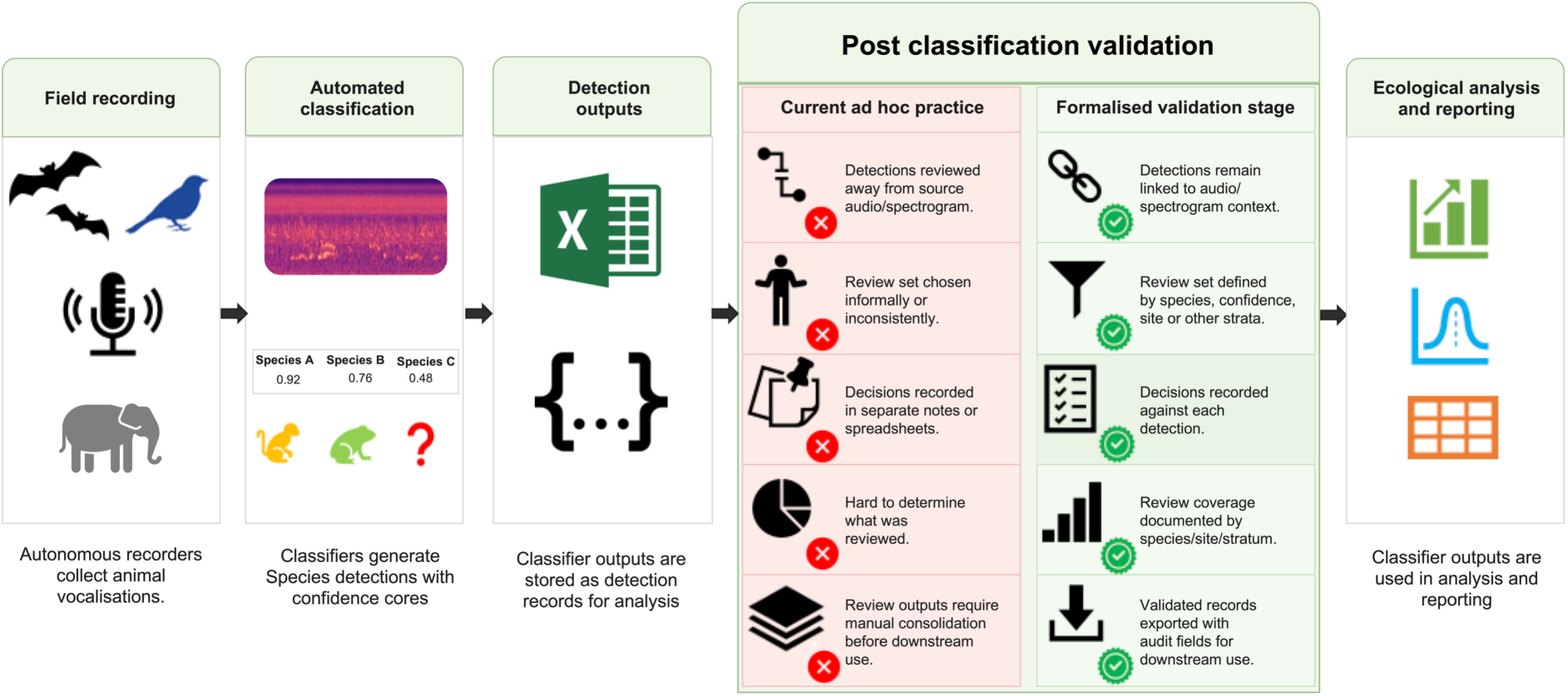
Conceptual workflow for passive acoustic monitoring showing the post-classification validation gap. Field recordings are processed by automated classifiers to produce detection outputs, which are then used in ecological analysis and reporting. The central panel expands the validation stage, contrasting illustrative examples of current ad hoc practice with a formalised workflow. The paired rows show how each approach handles links to source audio and spectrograms, review-set definition, decision recording, review coverage, and export of validated records.

The absence of a standardised validation step poses a number of challenges for how PAM outputs are used. Even when classifiers perform well on benchmark datasets, project-specific validation is important for quantifying error rates in the field, particularly where local soundscapes differ from training conditions (i.e., domain shift) (Metcalf *et al*., 2022). Ad hoc workflows frequently separate classifier predictions from the underlying audio, increasing manual handling and the scope for transcription or consolidation errors (Barré *et al*., 2019; López-Baucells *et al*., 2019; Malerba *et al*., 2026). Where validation is informal or weakly documented, false positives and false negatives may propagate unevenly into presence–absence summaries or occupancy analyses, risking biased or imprecise inferences (Chambert, Miller and Nichols, 2015; Russo and Voigt, 2016; Chambert *et al*., 2018; Barré *et al*., 2019; Katsis *et al*., 2025). Furthermore, the lack of a standardised post-classification validation step limits the creation of locally verified detections that can be used to calibrate decision thresholds or fine-tune classifiers under novel conditions (Lauha *et al*., 2022; Schiavo *et al*., 2025). Ad hoc workflows may also impose hidden costs, because time spent designing project-specific review processes, moving data between tools, reconciling files and documenting what was checked is time not spent reviewing detections. This may reduce review coverage for any resource-constrained team and could have disproportionate effects in monitoring contexts where staff time, funding, software access or specialist technical support are limited, including many low- and middle-income countries (Hahn, Bombaci and Wittemyer, 2022). Standardised validation workflows may therefore improve transparency and error control, while also helping limited review capacity to be used more efficiently.

Here, we introduce PAMalytics, an open-source, graphical user interface–based application designed to support post-classification validation as a standardised and reproducible stage of PAM workflows. PAMalytics was co-designed with practitioners and researchers from Conservation International Cambodia and Imperial College London to reflect the needs of applied monitoring and reporting programmes in different operational contexts. The software is classifier-agnostic, taking the tabular outputs of existing bioacoustic classifiers and structuring them into a validation workflow that reconnects each detection to its underlying audio and spectrogram within an interactive, browser-based environment. Pre-built adapters, which translate outputs from specific classifiers into the PAMalytics input format, are provided for widely used tools such as BirdNET and BatDetect2, while a general input format allows outputs from other classifiers to be incorporated. Users can review detections in a targeted way, choosing how much to review and where to focus effort, for example by prioritising particular species, sites, or the most and least confident detections. Validation decisions are recorded alongside the original classifier outputs, and review progress is tracked transparently, producing exportable detection tables. By providing these steps through a graphical, no-code interface, PAMalytics enables conservation practitioners to validate classifier outputs directly and supports a more traceable and auditable link between automated detections and downstream use in operational monitoring and assessment.

## 2 PAMalytics: Formalising post-classification validation

PAMalytics is designed to structure the post-classification validation stage by maintaining links between classifier predictions, source audio, contextual metadata, validation sampling decisions, reviewer judgements, and downstream analysis records. A user guide with installation and workflow instructions is available in the project repository.

The workflow is organised around six linked components: classifier outputs are standardised into detection-level validation records, users define a validation sampling strategy, detections are presented with spectrograms and audio playback for review, validation targets are set and tracked, different thresholds can be explored for sensitivity, and the validated detections are recorded and exported (Fig. 2).

**Figure 2:**
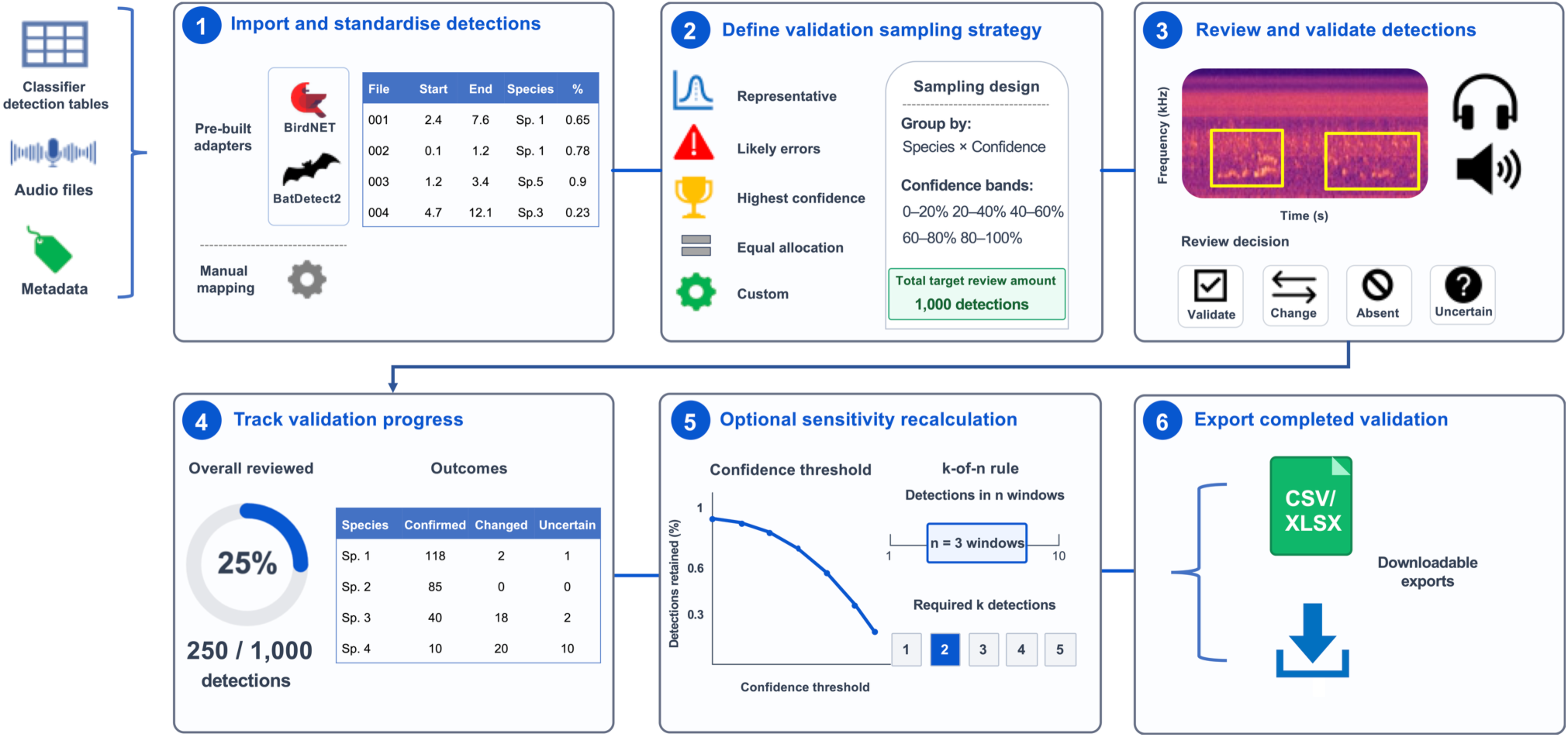
PAMalytics workflow conceptual diagram. The six panels show the main steps from importing and standardising classifier detections, audio files, and metadata; defining the validation sampling strategy; reviewing selected detections using spectrograms and audio playback; tracking review progress and outcomes; optionally recalculating derived summaries under post-classification rules; and exporting auditable records for downstream ecological analysis.

### 2.1 Standardising classifier outputs into detection-level validation records

PAMalytics reads in classifier outputs in tabular form, with each row representing one classifier detection and containing fields for file identity and path, detection start and end times, predicted labels or species, and classifier confidence. New validation fields are added to the detection table without overwriting the original classifier output meaning that original labels and confidence scores are retained for comparison. This structure preserves the relationship between the automated prediction, the source audio, and the human review outcome, while allowing downstream analyses to filter between unreviewed detections, confirmed detections, detections changed during validation, and detections highlighted for further review.

Currently PAMalytics contains built-in adapters for BirdNET and BatDetect2, as well as a guided manual mapping option which allows outputs from other classifiers to be incorporated. During project creation, classifier outputs are mapped to the common detection schema, and detections are linked to corresponding audio files. Metadata may be taken from the original detection tables or merged from separate files at the ingestion stage using shared identifiers. PAMalytics records the selected project metadata, column mappings, metadata joins, file path information, and validation status in a project manifest, allowing users to trace how the validation table was derived from the original classifier output and associated audio archive.

### 2.2 Defining a validation sampling strategy

Since reviewing every automated detection is often unfeasible, PAMalytics allows users to subset detections for validation according to project needs. A validation sampling wizard (Fig. 3) records the basis on which detections are selected, including the sampling strategy (Fig. 3a), grouping variable (e.g., species or site) (Fig. 3b), and target sample size (Fig. 3c).

**Figure 3:**
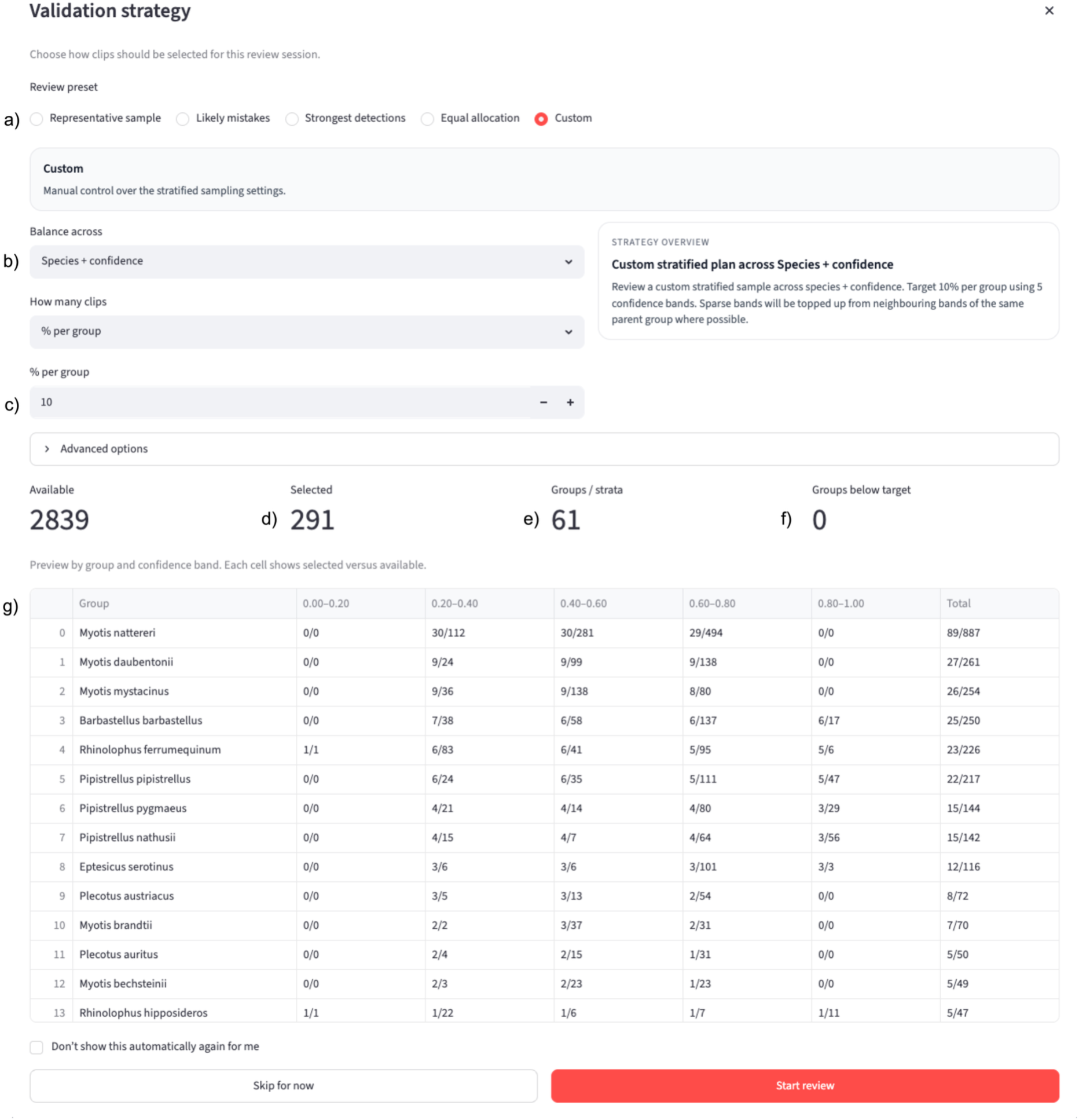
Validation strategy wizard screenshot. Users define the review preset (a), grouping structure (b), and target number of clips to review (c). In this example, a custom strategy selects 10% of detections per species, stratified across five classifier confidence bands from 0–20% to 80–100%. PAMalytics summarises the number of available detections, selected detections (d), groups or strata represented (e), and groups below target (f). The preview table (g) shows the number of selected and available detections for each species within each confidence band, together with the total selected and available detections per species.

Available strategies include representative sampling, which selects a random sample stratified by user-defined grouping variable; likely classifier mistakes, which prioritises low-confidence detections; strongest detections, which prioritises high-confidence detections and is useful when the goal is assessing site-level occupancy; equal allocation, which distributes the target sample evenly across confidence bands; and custom selection, which allows users to define their own stratified review set (Fig. 3a). Users can structure sampling by species, site, confidence band, or combinations of these variables.

Before a review set is created, PAMalytics displays the proposed allocation relative to the available detections. This includes the total target number of detections to be reviewed (Fig. 3d), the number of groups or strata represented (Fig. 3e), any groups below the requested target (Fig. 3f), as well as the number of selected and available detections per group (Fig. 3g). The resulting review set is stored with the user-defined sampling approach, allowing later summaries of validation coverage and outcomes to be interpreted in relation to the intended review design.

### 2.3 Validating detections with spectrograms and audio playback

In order to reduce the time spent navigating between different applications, the selected sample is passed through to the acoustic validation interface which combines the spectrogram view with audio playback and validation decision controls (Fig. 4). Detections from the same recording are displayed together on a shared spectrogram, with each selected detection annotated by its classifier confidence score (Fig. 4a). Reviewers can listen to the corresponding audio in the browser (Fig. 4b), can confirm the classifier predictions as correct, or revise the species label (Fig. 4c), and can additionally indicate that any of these decisions is uncertain (Fig. 4d). Uncertain records can then be revisited in a subsequent review. Where all displayed detections from the same recording are correct, reviewers can confirm them in a single step while retaining separate detection records (Fig. 4e). Review decisions are stored at detection level, linking each outcome to the classifier prediction and acoustic context used during validation.

**Figure 4:**
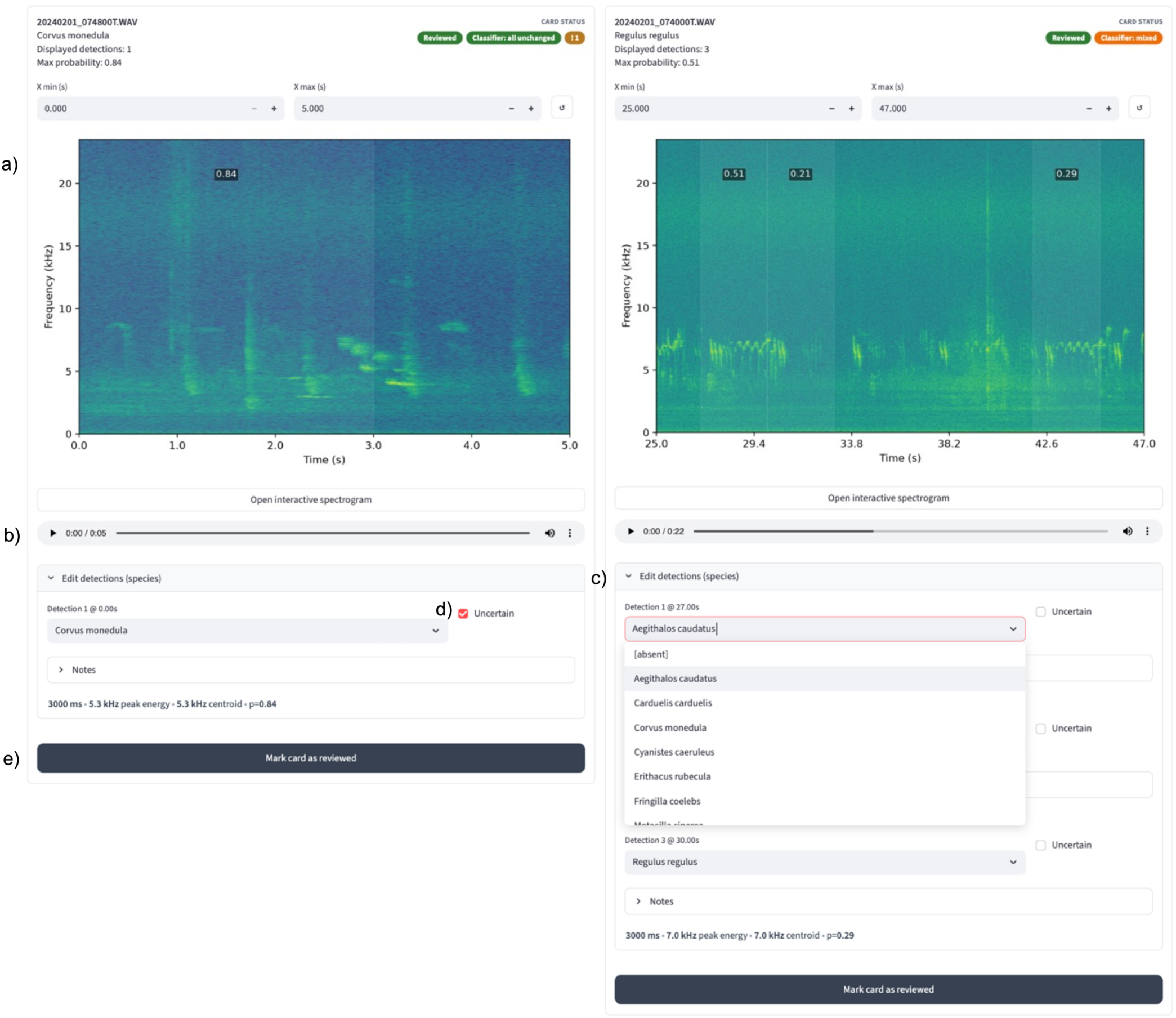
Screenshot of example validation screen. The screenshot shows two recording cards, each with a spectrogram panel (a), browser audio playback (b), per-detection label fields (c), uncertainty checkboxes (d), and card-level review controls (e).

### 2.4 Quantifying validation coverage and review outcomes

PAMalytics tracks validation status throughout the review process, allowing reviewers to see how much validation has been completed and how much remains. Summary tables report the number and proportion of detections reviewed, and the proportion confirmed, changed or flagged among reviewed detections, split by species and other grouping variables (Supplementary Fig. S1). These summaries also show how validation effort is distributed across the dataset, identify records or strata that remain unreviewed, and highlight species or groups where classifier predictions are frequently changed or marked as uncertain.

### 2.5 Applying optional post-classification rules and sensitivity analyses

Once a subset of detections has been validated, users can use the validation decisions to assess how the classifier performs under project-specific conditions and to decide how conservatively classifier outputs should be used. PAMalytics supports this by allowing users to adjust confidence thresholds and repeat-detection rules within the application, then compare how these settings affect the number of detections retained and the validation decisions associated with those detections. The validated detections retained at each threshold can then be used to compute performance measures such as precision, recall, and false-positive rate (Supplementary Fig. S2).

For example, a lower confidence threshold may retain more detections but include more false positives, while a higher threshold may retain fewer detections but a greater proportion of confirmed detections. This allows users to examine the trade-off between sample size and apparent error rate using the reviewed subset. When multiple detections of the same species occur within a recording, users can apply detection aggregation rules that group detections within a defined time window and retain detections only when there is sufficient supporting evidence from nearby detections. This avoids treating isolated detections as equally reliable and can reduce the influence of false positives, particularly for species with longer calling phases or repeated call structures (Supplementary Fig. S2).

### 2.6 Exporting auditable validation outputs

After validation, PAMalytics exports detection-level CSV and XLSX files containing the original classifier output, joined metadata, and validation fields. The export retains original species and presence labels alongside reviewer-assigned labels, validation state, uncertainty status, reviewer identity, and validation timestamp. Original and reviewed detections are therefore stored in the same record structure. The exported table can be incorporated into downstream scripts and reporting pipelines while preserving the information needed to condition analyses on review status, reviewer decision, uncertainty status, or classifier prediction. It can also provide validated labels for retraining or fine-tuning classifiers outside PAMalytics. This provides an auditable record of the validation stage and maintains the link between automated detections, validation decisions, and derived analytical outputs.

## 3 USE CASES

### 3.1 Conservation International: pileated gibbons in Cambodia

Pileated gibbons (*Hylobates pileatus*) are an endangered gibbon found in the central plain forests and southwestern mountain areas in Cambodia, south-east Thailand and south-west Laos (Brockelman, Timmins and Traeholt, 2020). The species shows strong associations with mature evergreen forest, and its densities decline with habitat disturbance and forest degradation, making it a valuable indicator of forest condition where direct biodiversity surveys are difficult (Phoonjampa *et al*., 2011). However, their arboreal and wide-ranging behaviour makes conventional survey methods logistically demanding (Neilson, Nijman and Nekaris, 2013). Passive acoustic monitoring has clear potential in this context because gibbons produce loud, stereotyped songs with consistent diurnal patterns and call-phrase durations of approximately 8–19 seconds (Traeholt *et al*., 2006). As part of a broader effort to develop standardised PAM workflows for biodiversity monitoring in Cambodia, the Conservation International (CI) Cambodia Biodiversity and Science team established a pilot programme for pileated gibbons in the Central Cardamom Mountains Landscape, southwestern Cambodia. Autonomous recording units were deployed across multiple forest sites to record morning periods over extended sampling windows, and the resulting audio was processed using a trained gibbon classifier to identify putative gibbon calls.

Before the introduction of PAMalytics, validation of automated gibbon detections within CI workflows relied on manual spot checking. Reviewers worked directly from classifier output tables, locating the corresponding audio files in local directories, opening each file in separate playback or visualisation software, and listening to the relevant sections to determine whether detections were correct. Because multiple detections were often associated with the same recording, this process was time-consuming and difficult to apply consistently across large datasets. The gibbon dataset was also imbalanced. Out of approximately 20,000 one-minute audio files recorded, only around 900 were predicted to contain a gibbon call, with large disparities in positive predictions across sites. This meant that a subset of the positive predictions could be validated to identify false positives, whereas validating the much larger set of negative predictions for missed calls was not feasible.

With PAMalytics, CI reviewers now conduct a preliminary review of positive detections based on ecological expectations, such as calls occurring outside typical calling periods, or on classifier confidence scores close to decision thresholds. This allows limited expert time to be directed towards detections that are most likely to contain errors, rather than relying on unstructured spot checking. Reviewers then subsample a defined target proportion of detections within each site and carry out validation directly within the application, using the integrated spectrogram and audio views. The time saved in the validation of positive calls also allows reviewers to use PAMalytics to examine candidate false negatives. For example, reviewers apply the detection aggregation feature to identify negative or low-confidence periods occurring near validated gibbon detections. These candidate missed-call periods are prioritised for additional review, allowing validation effort to address both false positives and plausible false negatives within the same workflow.

### 3.2 Imperial College London: Monitoring biodiversity co-benefits of large-scale restoration in the Brazilian Amazon

Tropical forest restoration is being implemented at increasingly large scales as a route to delivering both carbon and biodiversity outcomes (Strassburg et al., 2020; Edwards et al., 2021), yet robust evidence of biodiversity recovery - particularly for fauna - remains scarce relative to structural metrics such as canopy cover (Brancalion et al., 2025). As part of a programme monitoring the biodiversity co-benefits of large-scale restoration in the Brazilian Amazon, researchers at Imperial College London deployed passive acoustic monitoring across a network of restoration and reference sites in the state of Pará. Autonomous recorders (AudioMoth and Song Meter Mini units) were deployed at fixed sampling points distributed across former cattle-pasture sites undergoing native-forest restoration, alongside degraded baseline sites and old-growth forest reference controls, with recordings collected over repeated annual sampling windows. The resulting audio was processed with BirdNET to generate putative bird detections, producing detection tables substantially larger than could be reviewed in full. Therefore, a structured validation step was required before community-level metrics could be derived with confidence. Because the monitoring is intended to track recovery trajectories across many sites and years, and to feed into biodiversity reporting and credit-market contexts (Cord et al., 2025; Miller et al., 2025), the validation step needed to be both transparent and repeatable as well as efficient.

Before adopting PAMalytics, validation of BirdNET outputs followed an ad hoc, manual workflow. Reviewers worked directly from classifier output tables, manually subsetting detections in spreadsheets, locating the corresponding audio files by file path in local directories, and opening each file in separate audio software (e.g., Audacity) to listen to and inspect the relevant segment. Validation decisions were recorded in a separate spreadsheet, disconnected from the audio they referred to. This repeated movement between spreadsheet, file system, and playback software was slow, difficult to apply consistently across hundreds of species and thousands of detections and offered little transparency in recording how much of the dataset had been reviewed or how review effort was distributed across species, sites and confidence scores.

With PAMalytics, the same validation step is conducted within a single application. BirdNET detection tables are ingested and linked automatically to their source audio, so each detection can be reviewed in its spectrogram and audio context without manual file handling. The team applied a stratified validation design - retaining species with more than 50 detections and subsampling a fixed number per species across confidence-score bins - and PAMalytics supports this directly by allowing reviewers to filter to focal species and order detections by confidence score, ensuring even review coverage across the full confidence gradient so that the precision-confidence relationship can be estimated easily. Decisions are recorded per detection alongside the original classifier predictions rather than in a separate sheet, and review coverage is tracked transparently across species and sites. The exported table preserves both the original BirdNET predictions and the validation outcomes, feeding directly into the project’s species-specific confidence threshold calibration (Wood and Kahl, 2024) and downstream analyses. In this project, PAMalytics was used to process a dataset of 1,476,362 recordings to validate a target of 15,000 BirdNET detections spanning 300 species, supporting validation across restoration and reference sites. Using PAMalytics reduced the time required to validate a comparable set of detections from approximately 54.1 to 47.8 minutes per 100. Extrapolated across the full validation target, this equates to a reduction in reviewer time from approximately 135.2 hours under the manual approach to 119.5 hours using PAMalytics, a saving of 15.8 hours, alongside a clear and auditable record of review coverage. This estimate also does not include the time saved on the initial processing steps required for the manual method.

## 4 DISCUSSION

The increasing scale of passive acoustic monitoring and the routine use of automated classifiers have shifted PAM into operational monitoring and assessment (Aide *et al*., 2013; Browning *et al*., 2017). In this setting, large volumes of automated outputs frequently require quality assurance for robust reporting and analysis. PAMalytics addresses this downstream bottleneck by supporting post-classification validation as a distinct workflow stage. It links detections back to source audio and spectrograms, supports targeted review strategies, and records review outcomes alongside the original predictions. This combination helps make validation effort transparent and repeatable, while producing validated detection tables that can be passed into existing analytical pipelines without major restructuring.

The CI Cambodia and Imperial College London case studies show that PAMalytics increases the amount of validation that can be completed under realistic review constraints. In the CI workflow, this gain comes from directing scarce expert attention towards the gibbon detections most likely to affect interpretation of a sparse focal-species dataset, including plausible missed calls. In the Imperial workflow, it comes from removing the repeated spreadsheet, file-search, and audio-software steps that limited the pace of multi-species BirdNET validation. Across both cases, PAMalytics shifts reviewer time towards validation decisions and away from manual handling, allowing larger and more consistent review sets to be completed with fewer opportunities for transcription or consolidation error.

These examples point to a broader role for post-classification validation in PAM workflows. When classifier outputs are large, unevenly distributed, or strongly context-dependent, validation becomes a question of how review effort should be allocated across the dataset. Decisions about which detections to check, how to balance effort across species, sites or confidence scores, and how to interpret reviewed records alongside unreviewed predictions all affect the reliability of downstream summaries. PAMalytics makes these decisions explicit by offering several ways to define review sets, including representative sampling, confidence-based review, equal allocation, and custom selection. The application does not recommend a universally preferred sampling strategy, so the appropriate choice remains study-specific and should be guided by the monitoring objective, expected error structure, available reviewer capacity, and intended downstream analysis.

The significance of this workflow also depends on whether practitioners can implement it in practice. A central benefit of PAMalytics is that it provides a no-code, local browser-based graphical interface for post-classification validation. This allows conservation practitioners with ecological or taxonomic expertise to perform structured validation directly, even where access to data-science support is limited. The software still depends on reviewer expertise and an appropriate review design, but it reduces the technical barrier to applying those elements consistently.

This has practical importance where biodiversity monitoring capacity is uneven. Skills, cost, and data-management constraints are widely recognised barriers to the uptake of conservation technologies in the field, and tools that reduce coding requirements, manual handling, and workflow fragmentation can help address these constraints (Hahn, Bombaci and Wittemyer, 2022). In organisations with limited data-science capacity, including many in low- and middle-income countries, structured no-code validation applications can make automated acoustic outputs easier to review, document and use in local decision-making and reporting. PAMalytics therefore contributes both a software implementation and a practical framework for making post-classification validation more transparent, reproducible, and accessible in applied monitoring contexts.

## Acknowledgements

This research was supported by a Natural Environment Research Council grant to AP [grant number NE /S007229/1]

## REFERENCES

Aide, T.M. et al. (2013) ‘Real-time bioacoustics monitoring and automated species identification’, PeerJ, 1, p. e103. Available at: 10.7717/peerj.103.

Aodha, O.M. et al. (2022) ‘Towards a General Approach for Bat Echolocation Detection and Classification’. bioRxiv, p. 2022.12.14.520490. Available at: 10.1101/2022.12.14.520490.

Barré, K. et al. (2019) ‘Accounting for automated identification errors in acoustic surveys’, Methods in Ecology and Evolution, 10(8), pp. 1171–1188. Available at: 10.1111/2041-210X.13198.

Brancalion, P.H.S. et al. (2025) ‘Moving biodiversity from an afterthought to a key outcome of forest restoration’, Nature Reviews Biodiversity, 1(4), pp. 248–261. Available at: 10.1038/s44358-025-00032-1.

Brockelman, W., Timmins, T. and Traeholt, C. (2020) ‘Hylobates pileatus. The IUCN Red List of Threatened Species 2020: e.T10552A17966665.’, IUCN Red List of Threatened Species [Preprint]. Available at: 10.2305/.

Browning, E. et al. (2017) ‘Passive acoustic monitoring in ecology and conservation.’

Chambert, T. et al. (2018) ‘Two-species occupancy modelling accounting for species misidentification and non-detection’, Methods in Ecology and Evolution, 9(6), pp. 1468–1477. Available at: 10.1111/2041-210X.12985.

Chambert, T., Miller, D.A.W. and Nichols, J.D. (2015) ‘Modeling false positive detections in species occurrence data under different study designs’, Ecology, 96(2), pp. 332–339. Available at: 10.1890/14-1507.1.

Cord, A.F. et al. (2025) ‘Leveraging passive acoustic monitoring for result-based agri-environmental schemes: Opportunities, challenges and next steps’, Biological Conservation, 305, p. 111042. Available at: 10.1016/j.biocon.2025.111042.

Cowans, A. et al. (2024) ‘Improving the integration of artificial intelligence into existing ecological inference workflows’, Methods in Ecology and Evolution, pp. 2041–210X.14485. Available at: 10.1111/2041-210X.14485.

Edwards, D.P. et al. (2021) ‘Upscaling tropical restoration to deliver environmental benefits and socially equitable outcomes’, Current Biology, 31(19), pp. R1326–R1341. Available at: 10.1016/j.cub.2021.08.058.

Gibb, R. et al. (2019) ‘Emerging opportunities and challenges for passive acoustics in ecological assessment and monitoring’, Methods in Ecology and Evolution, 10(2), pp. 169–185. Available at: 10.1111/2041-210X.13101.

Gillespie, D. et al. (2009) ‘PAMGUARD: Semiautomated, open source software for real-time acoustic detection and localization of cetaceans.’, The Journal of the Acoustical Society of America, 125(4_Supplement), p. 2547. Available at: 10.1121/1.4808713.

Hahn, N.R., Bombaci, S.P. and Wittemyer, G. (2022) ‘Identifying conservation technology needs, barriers, and opportunities’, Scientific Reports, 12(1), p. 4802. Available at: 10.1038/s41598-022-08330-w.

Kahl, S. et al. (2021) ‘BirdNET: A deep learning solution for avian diversity monitoring’, Ecological Informatics, 61, p. 101236. Available at: 10.1016/j.ecoinf.2021.101236.

Katsis, L.K.D. et al. (2025) ‘A comparison of statistical methods for deriving occupancy estimates from machine learning outputs’, Scientific Reports, 15(1), p. 14700. Available at: 10.1038/s41598-025-95207-3.

Lauha, P. et al. (2022) ‘Domain-specific neural networks improve automated bird sound recognition already with small amount of local data’, Methods in Ecology and Evolution, 13(12), pp. 2799–2810. Available at: 10.1111/2041-210X.14003.

López-Baucells, A. et al. (2019) ‘Stronger together: Combining automated classifiers with manual post-validation optimizes the workload vs reliability trade-off of species identification in bat acoustic surveys’, Ecological Informatics, 49, pp. 45–53. Available at: 10.1016/j.ecoinf.2018.11.004.

Malerba, M.E. et al. (2026) ‘Overcoming software bottlenecks for scalable passive acoustic monitoring: insights from a global expert assessment’. bioRxiv, p. 2026.03.30.715176. Available at: 10.64898/2026.03.30.715176.

Marques, T.A. et al. (2013) ‘Estimating animal population density using passive acoustics’, Biological Reviews, 88(2), pp. 287–309. Available at: 10.1111/brv.12001.

Martínez Balvanera, S., et al. (2025) ‘Whombat: An open-source audio annotation tool for machine learning assisted bioacoustics’, Methods in Ecology and Evolution, 16(1), pp. 19–28. Available at: 10.1111/2041-210X.14468.

Metcalf, O.C. et al. (2022) ‘Detecting and reducing heterogeneity of error in acoustic classification’, Methods in Ecology and Evolution, 13(11), pp. 2559–2571. Available at: 10.1111/2041-210X.13967.

Miller, B.L. et al. (2025) ‘Corporate Biodiversity Reporting Can Be Scaled With AI and Earth Observation—But Will Miss the Point Without Guidance From Conservation Scientists’, Conservation Letters, 18(5), p. e13153. Available at: 10.1111/conl.13153.

NatureScot (2021) Bats and onshore wind turbines - survey, assessment and mitigation. Available at: https://www.nature.scot/doc/bats-and-onshore-wind-turbines-survey-assessment-and-mitigation (Accessed: 16 December 2025).

Neilson, E., Nijman, V. and Nekaris, K.A.I. (2013) ‘Conservation Assessments of Arboreal Mammals in Difficult Terrain: Occupancy Modeling of Pileated Gibbons (Hylobates pileatus)’, International Journal of Primatology, 34(4), pp. 823–835. Available at: 10.1007/s10764-013-9688-6.

Phoonjampa, R. et al. (2011) Pileated gibbon density in relation to habitat characteristics and post-logging forest recovery. Available at: https://repository.li.mahidol.ac.th/entities/publication/5e24635c-6a56-4851-ad6c-84b16baf1a31 (Accessed: 30 December 2025).

Russo, D. and Voigt, C.C. (2016) ‘The use of automated identification of bat echolocation calls in acoustic monitoring: A cautionary note for a sound analysis’, Ecological Indicators, 66, pp. 598–602. Available at: 10.1016/j.ecolind.2016.02.036.

Schiavo, G. et al. (2025) ‘Fine-Tuning BirdNET for the Automatic Ecoacoustic Monitoring of Bird Species in the Italian Alpine Forests’, Information, 16(8). Available at: 10.3390/info16080628.

Stowell, D. (2022) ‘Computational bioacoustics with deep learning: a review and roadmap’, PeerJ, 10, p. e13152. Available at: 10.7717/peerj.13152.

Strassburg, B.B.N. et al. (2020) ‘Global priority areas for ecosystem restoration’, Nature, 586(7831), pp. 724–729. Available at: 10.1038/s41586-020-2784-9.

Sugai, L.S.M. et al. (2019) ‘Terrestrial Passive Acoustic Monitoring: Review and Perspectives’, BioScience, 69(1), pp. 15–25. Available at: 10.1093/biosci/biy147.

Traeholt, C. et al. (2006) ‘Song Activity of the Pileated Gibbon, Hylobates pileatus, in Cambodia’, Primate Conservation, 2006(21), pp. 139–144. Available at: 10.1896/0898-6207.21.1.139.

Wood, C.M. and Kahl, S. (2024) ‘Guidelines for appropriate use of BirdNET scores and other detector outputs’, Journal of Ornithology, 165(3), pp. 777–782. Available at: 10.1007/s10336-024-02144-5.

Wright, W.J. et al. (2020) ‘Modelling misclassification in multi-species acoustic data when estimating occupancy and relative activity’, Methods in Ecology and Evolution, 11(1), pp. 71–81. Available at: 10.1111/2041-210X.13315.

