## Supplementary Fig for "PAMalytics: a no-code application for structured validation of bioacoustic detections"

**Supplementary Information**

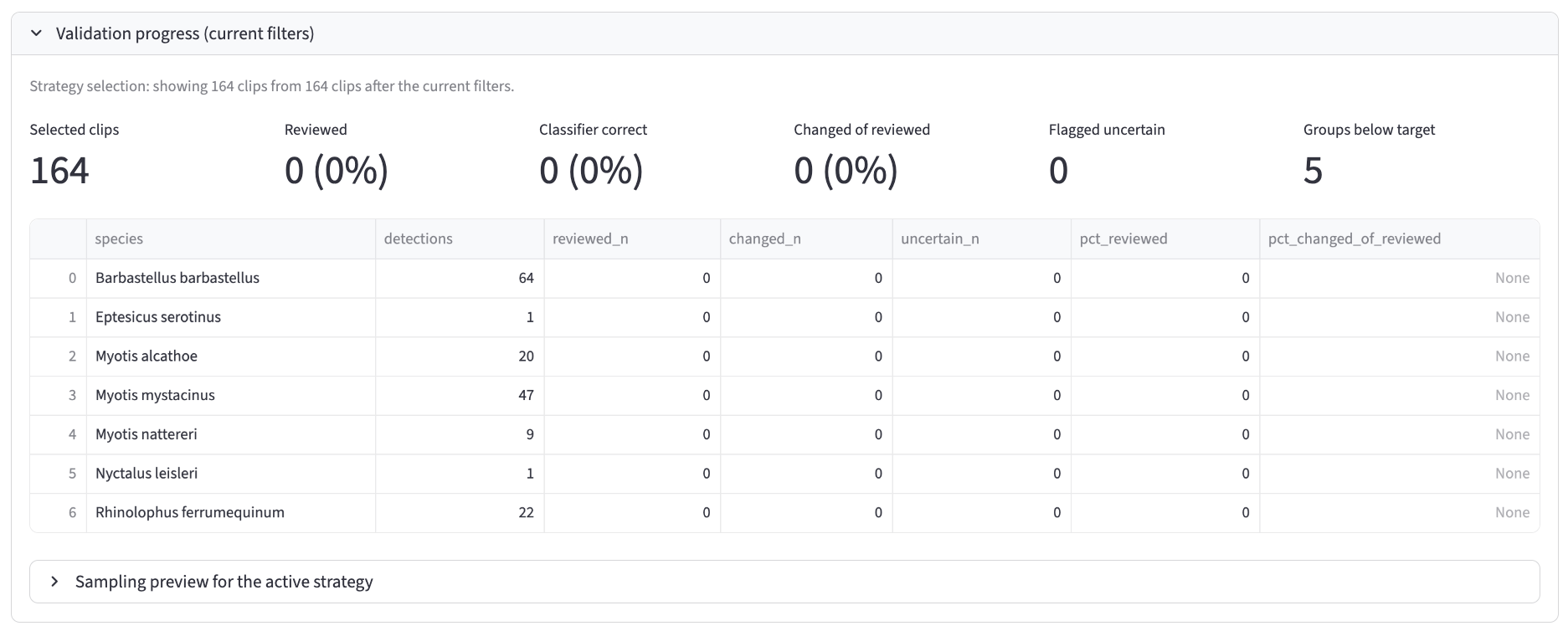

**Supplementary Figure S1: Validation progress tracker screenshot**

*Table and summary displaying the total proportion of detections that have been reviewed and the proportions that are correct, uncertain or require reviewer changes both overall and per grouping variable (species or site).*

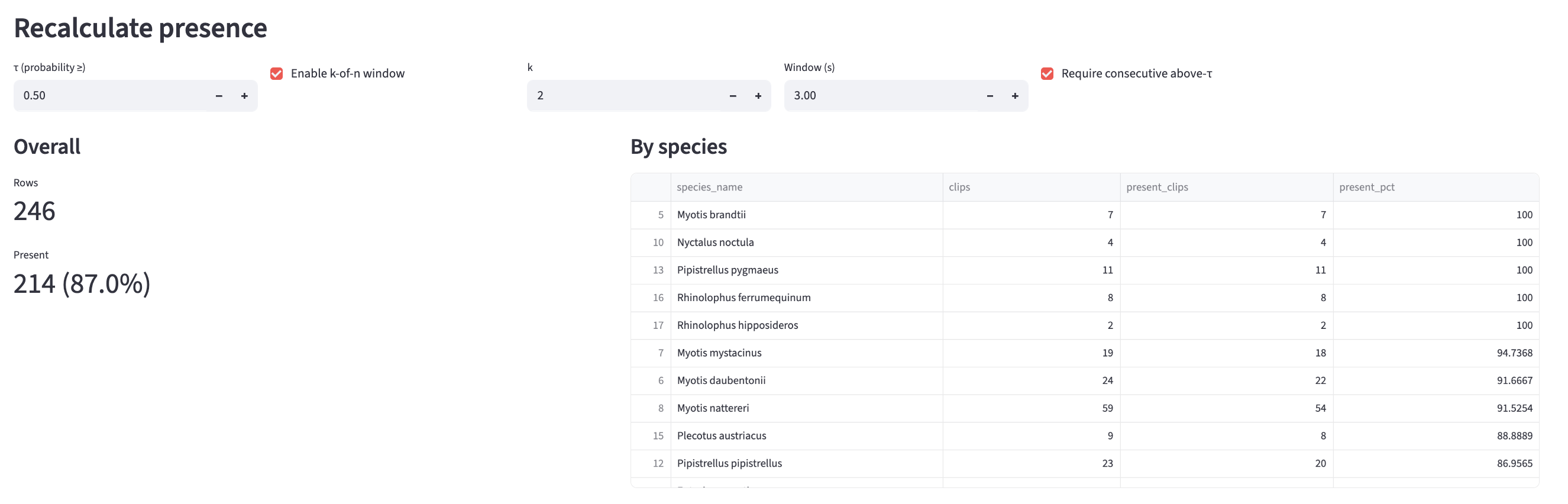

**Supplementary Figure S2: Recalculate page screenshot**

*Users can change the presence threshold (shown here as 0.5) and/or update the presence rule to enable a k-of-n approach (k detections in n windows, shown here as 2 detections in 3 windows). This will recalculate the summary totals.*
